# Control of subunit stoichiometry in single-chain MspA nanopores

**DOI:** 10.1101/2021.09.25.461773

**Authors:** Mikhail Pavlenok, Luning Yu, Dominik Herrmann, Meni Wanunu, Michael Niederweis

## Abstract

Transmembrane protein channels enable fast and highly sensitive electrical detection of single molecules. Nanopore sequencing of DNA was achieved using an engineered *Mycobacterium smegmatis* porin A (MspA) in combination with a motor enzyme. Due to its favorable channel geometry, the octameric MspA pore exhibits the highest current level as compared to other pore proteins. To date, MspA is the only protein nanopore with a published record of DNA sequencing. While widely used in commercial devices, nanopore sequencing of DNA suffers from significant base-calling errors due to stochastic events of the complex DNA-motor-pore combination and the contribution of up to five nucleotides to the signal at each position. Asymmetric mutations within subunits of the channel protein offer an enormous potential to improve nucleotide resolution and sequencing accuracy. However, random subunit assembly does not allow control of the channel composition of MspA and other oligomeric protein pores. In this study, we showed that it is feasible to convert octameric MspA into a single-chain pore by connecting eight subunits using peptide linkers. We constructed single-chain MspA trimers, pentamers, hexamers and heptamers to demonstrate that it is feasible to alter the subunit stoichiometry and the MspA pore diameter. All single-chain MspA proteins formed functional channels in lipid bilayer experiments. Importantly, we demonstrated that single-chain MspA discriminated all four nucleotides identical to MspA produced from monomers. Thus, single-chain MspA constitutes a new milestone in its development and adaptation as a biosensor for DNA sequencing and many other applications.

**STATEMENT OF SIGNFICANCE:** Nanopore sequencing of DNA is a fast and cheap technology that uniquely delivers multi-kilobase reads. It is currently used world-wide in many applications such as genome sequencing, epigenetics, and surveillance of viral and bacterial pathogens and has started to revolutionize human lives in medicine, agriculture and environmental studies. However, the high base-calling error rates prevent nanopore DNA sequencing from reaching its full potential. In this study, we converted octameric MspA into a single-chain pore enabling asymmetric mutations to fine-tune the pore geometry and chemistry and address the shortcomings of nanopores. Thus, single-chain MspA constitutes a new milestone in its development and adaptation as a biosensor for DNA sequencing and many other applications.

## INTRODUCTION

Protein pores have attracted an enormous interest as biosensors as they enable fast and highly sensitive detection of analytes by recording current traces. This concept of stochastic sensing was pioneered using bacterial and fungal pores (1,2) and has been applied to a wide variety of small molecules and polymers (3). In particular, DNA sequencing with pore proteins is a promising technology due to its low cost, fast processing, minimal sample amount and lack of sample amplification (4). While the concept of nanopore DNA sequencing was developed using the staphylococcal α-hemolysin pore (5), it was demonstrated to be a feasible technology using the *Mycobacterium smegmatis* porin A (MspA) (6). MspA is the most stable of the known pore proteins resisting denaturation by 2% SDS at 100 °C (7) and is functional in many different membrane-like environments (8-10), an important feature for technical applications (3). We first engineered MspA to translocate DNA (11), showing that MspA can distinguish the four nucleobases (12) and then achieved nanopore sequencing of DNA in combination with a motor enzyme (13). Several other channel proteins have also been used for nucleotide sensing including α–hemolysin (14), CsgG (15), ClyA (16), FraC (17) and aerolysin (18). MspA exhibits the highest current level compared to α–hemolysin, CsgG and aerolysin, likely due to its short channel constriction and its wide vestibule at the pore entrance (19), increasing the signal-to-noise ratio. MspA is the only protein nanopore with a published record of DNA sequencing (13,20). Mutants of the CsgG nanopore are apparently being used in the commercially available MinION devices (21), but no molecular details of the performance of CsgG in DNA sequencing experiments have been published to our knowledge.

One of the main problems of nanopore sequencing is the very fast translocation of DNA (22). Control of the DNA translocation rate is currently achieved by DNA-processing enzymes, but this adds complexicity and stochastic signals from the motor protein, decreasing the signal-to-noise ratio (13,23). In addition, the residual current of single-stranded DNA passing through the pore is determined by four to five nucleotides at each position (13). These and other limitations result in raw base calling errors up to 12% for MspA (24). To our knowledge, raw base calling errors are not published for any other nanopore. Many of these challenges could, in principle, be addressed by protein engineering. However, the oligomeric nature of MspA and of other protein nanopores comprising seven to twelve subunits, severely limits these efforts (25). Asymmetric mutations are required to efficiently improve nucleotide recognition, to better control DNA capture and translocation rates and to control the path of DNA translocation. While first steps in this direction have been made by electrophoretically separating permutations of a mixture of two distinct hemolysin subunits after chemical modifications (26), this process is tedious, difficult to scale and is not usable for a combination of mutations in different subunits. This can only be effectively achieved by synthesizing one gene which encodes all subunits. We demonstrated that this is feasible for MspA because its C-terminus is close to the N-terminus of the next subunit (Fig. 1A) and showed that two covalently connected subunits indeed assemble to functional MspA pores (25). However, obtaining pure single-chain MspA, in which all eight subunits are linked (Fig. 1B), has been elusive for almost a decade.

**Fig. 1.**
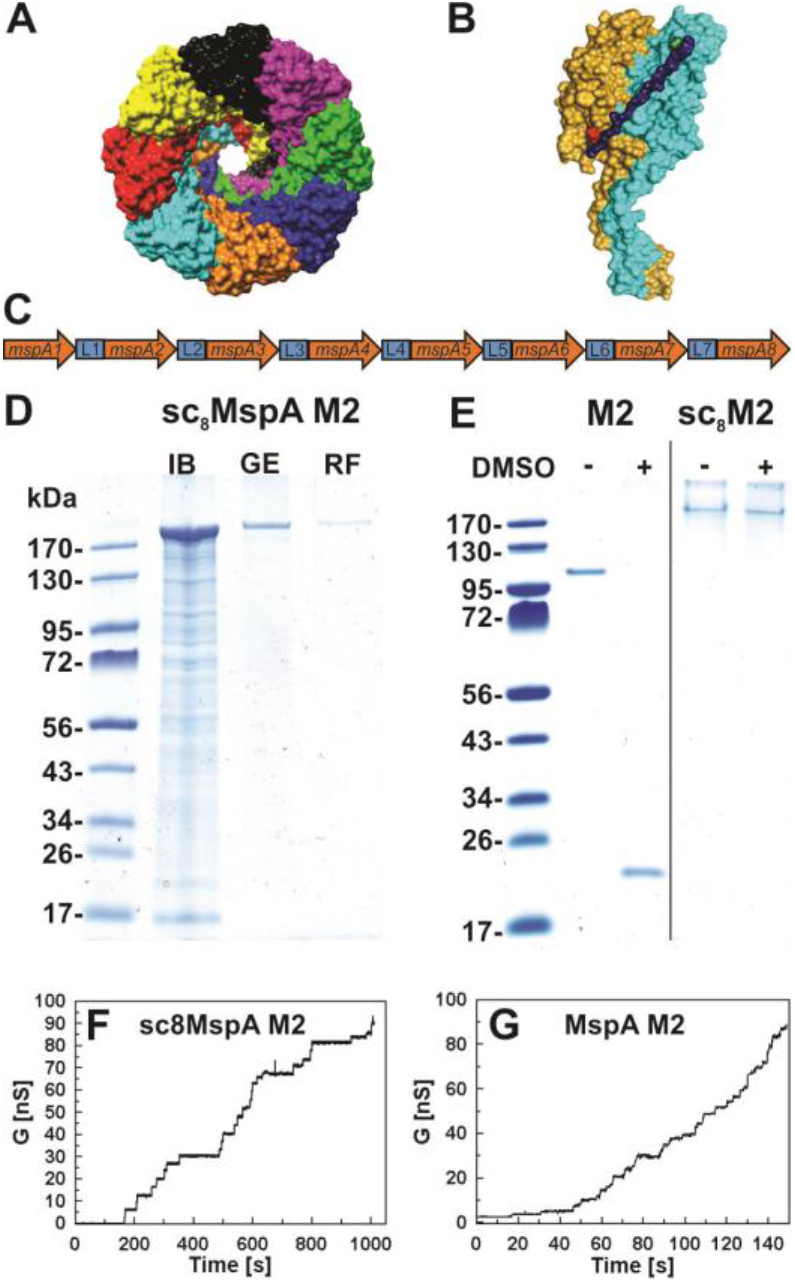
Design, production and purification of single-chain MspA. **(A)** Structure of wt MspA (PDB#: 1UUN) with eight monomers in different colors (top view). **(B)** Model of the covalent peptide linker between two MspA subunits in single-chain MspA (side view). The linker connects the C-terminus (red) of one subunit to the N-terminus (blue) of the adjacent subunit. Models in A and B were prepared using Chimera 11. **(C)** The single-chain *mspA* gene. Numbered arrows represent translationally fused *mspA m2* genes. Blue boxes with numbers show the linker regions connecting adjacent *mspA m2* genes.. The scheme is not to scale. **(D)** Purification of sc_8_MspA M2. Samples were loaded onto 10% polyacrylamide gel followed by staining with Coomassie. Lanes: IB - inclusion bodies purified from Omp8 *E*.*coli* solubilized in 8 M urea; GE - sample after gel extraction procedure; RF – refolded sample after dialysis. Numbers on the left are molecular weights of the marker in kDa. **(E)** Denaturation of MspA M2 and s_8_MspA M2. Coomassie-stained 8% polyacrylamide gel of untreated (-) and samples boiled in 80% (v/v) DMSO (+) to denature proteins. Lanes: M2 – octameric MspA M2; sc_8_M2 – sc_8_MspA M2 with eight covalently linked subunits. 1 µg of protein was loaded on each lane. Current trace of sc_8_MspA M2 **(F)** and octameric MspA M2 **(G)** in a diphytanoylphosphatidylcholine (DPhPC) membrane with an aperture diameter of 1 mm in a Montal-Mueller system. The applied potential was −10 mV, the electrolyte was 1 M KCl, 10 mM Hepes (pH 7.5).

In this study, we show that it is feasible to convert octameric MspA into a monomeric pore enabling asymmetric mutations to optimize the pore properties for specific applications. As proof-of-principle, we also constructed and purified single-chain MspA trimers, pentamers, hexamers, and heptamers, highlighting the possibility to construct pores with different channel diameters by controlling their subunit stoichiometry. All single-chain MspA proteins formed functional channels in lipid bilayer experiments. Importantly, we demonstrated that full-length single-chain MspA discriminated all four nucleotides in a manner identical to MspA produced from monomers. Single-chain MspA constitutes a new milestone in its development and adaptation as a nanosensor for DNA sequencing and many other applications.

## MATERIAL AND METHODS

**Chemicals and Reagents, bacterial strains, media and growth conditions and standard procedures** are described in the ***SI Appendix***

### Plasmid construction

Standard molecular biology methods were used for plasmid construction. Single-chain MspA genes were ordered from GenScript (Piscataway, NJ). Description of construction of single-chain MspA M2 and single-chain MspA PN1 plasmids are in the ***SI Appendix***.

### Purification of MspA octamers

Purification of octameric MspA from *M*.*smegmatis* was performed as described elsewhere (27) and is summarized in the ***SI Appendix***.

### Expression and purification of single-chain MspA proteins

Single-chain MspA protein were purified from *E*.*coli* BL21(DE3)omp8 strain (28). After induction with 1 mM IPTG, cells were harvested and inclusion bodies were purified and solubilized in 8 M urea. Affinity purification or gel excision were used to further purify single-chain proteins. The proteins were then refolded by dialysis against buffer containing 0.5% OPOE detergent. For detailed purification procedures of single-chain MspA proteins see ***SI Appendix***.

### Lipid bilayer experiments and single-chennel hairpin experiments

Lipid bilayer experiments to check pore formation by MspA proteins were performed in diphytanoylphosphatidylcholine artificial membranes in a custom made bilayer apparatus as previously described (27) and summarized in ***SI Appendix***. Single-channel recodings and hairpin experiments were done in SU-8 wedge-on-pillar aperture and poly(1,2-butadiene)-b-poly(ethylene oxide) copolymer was used to form a bilayer as described (10). Data were collected at 250 kHz sampling rate and digitized. For details see ***SI Appendix***.

## RESULTS

### Purification and characterization of single-chain MspA

Eight MspA subunits assemble in the outer membrane of *Mycobacterium smegmatis* to produce a central water-filled channel (Fig. 1A). We have shown previously that the C-terminus of a monomer can be connected to the N-terminus of a neighboring subunit by a 17-mer peptide linker due to the close proximity of the termini (25). Here, we used the same approach to synthesize a single-chain *mspA m2* (*scmspA m2*) gene where eight *mspA m2* genes encoding the mutations D90N/D91N/D93N/D118R/D134K/D139R, which enable efficient DNA capture and translocation (11), are connected by DNA fragments encoding (GGGGS)_3_ peptide linkers (Fig. 1B; Table S1). Each gene is flanked by unique restriction sites to enable specific modifications of each MspA subunit (Table S1). We first tried to produce scMspA M2 protein in the porin quadruple mutant *M. smegmatis* ML712, which lacks all four *msp* genes (29). However, all *scmspA m2* genes in clones producing MspA protein were scrambled and protein quantities were very low (not shown). Then, we used *E. coli* to produce scMspA M2 protein. To reduce the probability of homologous recombination between the individual *mspA* genes, we altered every second codon in the *mspA* subunit genes 2 through 8 to generate DNA sequence differences without altering the amino acid sequence. In addition, the *scmspA m2* DNA sequence was altered for optimal expression in *E. coli* and the signal peptide was removed for cytoplasmic expression of MspA. We expressed the synthetic *scmspA m2* gene under the control of the T7 promoter using the plasmid pML3216 (Fig. S1) in the BL21(DE3)omp8 strain lacking the three major porins of *E. coli* (28) to avoid contamination with endogenous porins. MspA protein (166 kDa theoretical mass) was purified from inclusion bodies by gel electrophoresis and excision of a protein band of 170 kDa and refolded in a buffer contgaining the detergent octyl polyoxyethylene (OPOE) (Fig. 1D). The protein yield was 12 µg of purified sc_8_MspA M2 per single preparation from four polyacrylamide gels (approximately 190 µg per liter of *E. coli* culture).

Octameric MspA is an extremely stable protein that does not denature after boiling for 10 min in 2% SDS and other harsh denaturing conditions (7). To dissociate octameric MspA into its subunits we boiled the purified proteins in 80% (v/v) dimethylsulfoxide (DMSO) (25). Only octameric MspA M2 dissociated into monomers, while sc_8_MspA M2 was stable demonstrating that all eight subunits are covalently linked and that we purified full-length scMspA (Fig. 1E). To examine whether sc_8_MspA M2 forms functional channels we performed lipid bilayer experiments in diphytanoyl phosphatidylcholine (DPhPC) membranes of 1 mm in diameter in a Montal-Mueller setup (25). Addition of both octameric MspA M2 purified from *M. smegmatis* (Fig. 1G) and recombinant sc_8_MspA M2 protein to the membranes resulted in a step-wise current increase indicative of channel insertions (Fig. 1F), while no channel activity was observed when only detergent-containing buffer was added (not shown). Analysis of the single channel insertions revealed similar conductance values ranging from 0.5 to 6 nS for octameric MspA M2 and from 0.2 to 4.5 nS for single chain MspA (Fig. S4). These wide ranges of single channel conductances were also observed for other octameric MspA proteins such as wt MspA and MspA M1 (Fig. S4) and seem to be an intrinsic feature of the MspA pore. Taken together, these experiments demonstrate that sc_8_MspA M2 forms functional pores and show that it is feasible to convert an oligomeric pore protein into a functional monomeric protein enabling asymmetric mutations to improve channel properties.

### Single-chain MspA variants with altered subunit stoichiometries form functional channels

To highlight the advantages of scMspA in tailoring the pore for specific applications, we aimed at changing its channel diameter as one of the most important properties determining the interactions of the pore constriction with the analyte. Previously, the constriction diameter could only be altered by mutations of individual amino acids in the protein monomer, which also changes the chemical nature of the interactions with the analyte. A monomeric pore consisting of identical subunits enables for the first time to vary the channel diameter by altering the subunit stoichiometry. To this end, we designed scMspA variants with three, five, six, seven, and eight subunits based on MspA M2 with an additional P97F mutation (Table S5). This additional mutation was chosen because octameric MspA PN1 has a well defined channel distribution with a peak at 1.2 nS (Fig. S5A) in contrast to all other examined octameric MspA pores (Fig. S4). The corresponding proteins were named sc_x_MspA PN1 with “x” numbers of covalently linked subunits and were purified from *E. coli* BL21(DE3)omp8 as described above. The target proteins accounted for approximately 10% of proteins in the whole-cell lysate as determined by quantitative image analysis of protein gels (Fig. S3). The apparent molecular masses for sc_3_MspA PN1, sc_5_MspA PN1, sc_6_MspA PN1, and sc_7_MspA PN1 proteins were 60 kDa, 110 kDa, 120 kDa, and 140 kDa, respectively (Fig. 2A), and were in a good agreement with the theoretical molecular masses. To examine whether the scMspA PN1 subunits are covalently linked, the proteins were denatured by boiling in 80% (v/v) DMSO. DMSO treatment dissociated octameric MspA M2 protein into its 20 kDa monomers, while all scMspA PN1 proteins were stable under these conditions (Fig. 2A) demonstrating that the subunits of these proteins are indeed covalently linked. To examine the channel-forming properties of scMspA PN1 pores we performed planar lipid bilayer experiments as previously described (29). In these experiments addition of 70 – 840 ng of refolded proteins to the membrane resulted in a step-wise current increase indicative of channel insertions into lipid membranes (Fig. 2B-F). The channel insertion frequencies were similar for all scMspA PN1 proteins. The scMspA PN1 pores showed conductance values ranging between 0.5 to 5 nS (Fig. S5) similar to sc_8_MspA M2 and octameric MspA proteins (Fig. S4). Interestingly, the distribution peaks shifted to larger single channel conductances for the sc_3_MspA PN1, sc_5_MspA PN1 and sc_7_MspA PN1 variants (Figs. S5), indicating the presence of pores with more than eight subunits. Taken together, these experiments demonstrated that it is feasible to construct MspA pores with altered stoichiometries using covalently linked subunits. Thus, the single-chain concept enables the design of MspA pores with constriction zone diameters optimized for translocating different substrates, e.g. in nucleic acid and polypeptide sequencing.

**Fig. 2.**
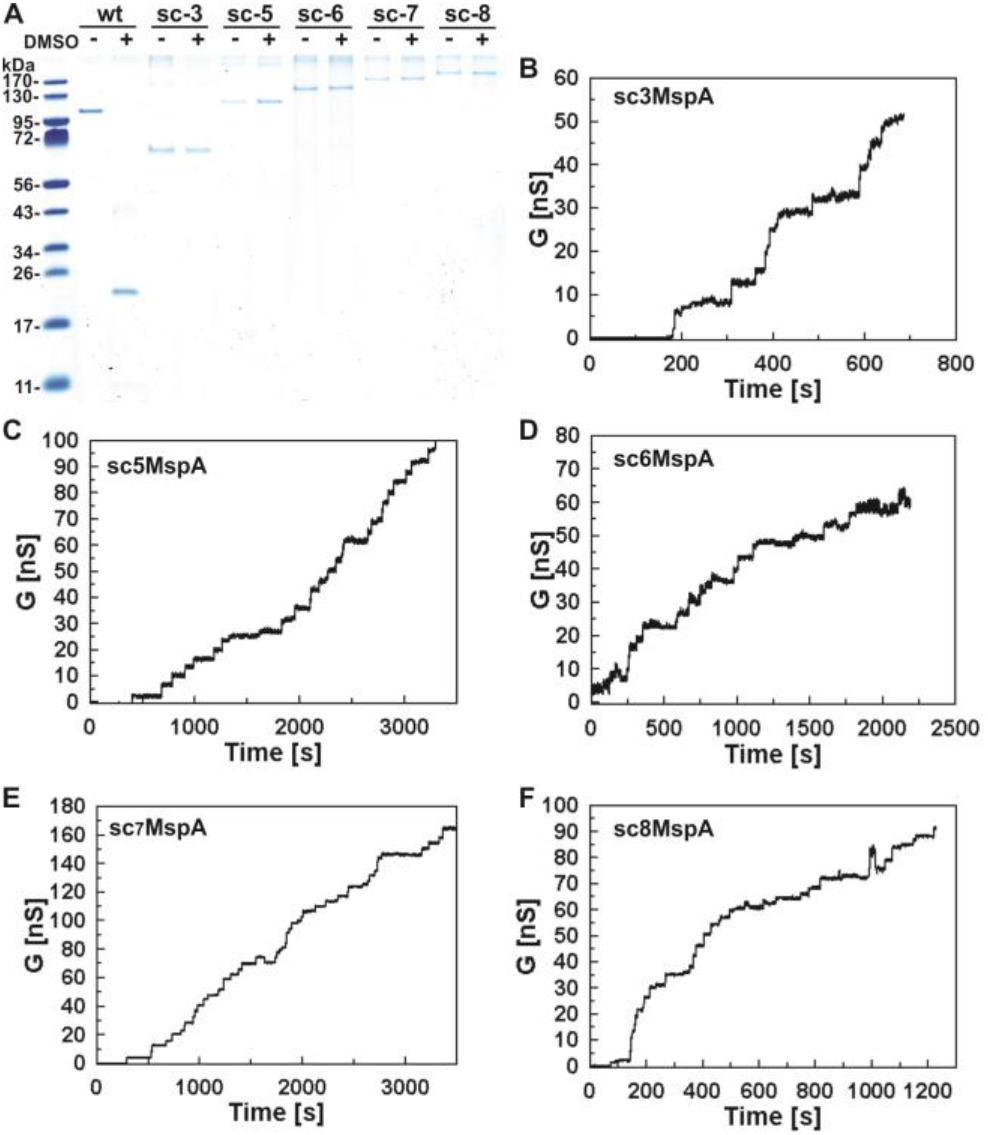
Channel activity of MspA pores with different subunit stoichiometries. **(A)**. Coomassie-stained 8% polyacrylamide gel with different single-chain MspA PN1 constructs. Lanes: wt – octameric MspA M2; sc-3, −5, −6, −7, −8 – single-chain MspA PN1 with three, five, six, seven, eight covalently linked subunits. Octameric MspA M2 and proteins with different subunit composition were boiled in 80% DMSO (+) and compared to untreated protein (-). Note that the 20 kDa band is monomeric subunit of MspA. Equal amounts (2 µg) of each sample were loaded on the gel. **(B - F)** Current traces of single-channel conductances of scMspA PN1 constructs. Membrane currents were recorded in 1 M KCl, 10 mM HEPES, pH 7.4 electrolyte at −10 mV applied potential. Diameter of the diphytanoyl phosphatidylcholine (DPhPC) bilayer was approximately 1 mm. **(B)** sc_3_MspA PN1: 95 insertion events from 4 membranes; **(C)** sc_5_MsA PN1: 221 insertions from 7 membranes. **(D)** sc_6_MspA PN1: 463 insertion events from 7 different membranes. **(E)** sc_7_MspA PN1: 62 insertions from 5 membranes; **(F)** sc_8_MspA PN1: 150 insertions from 6 membranes.

### Improvement of the purification of recombinant single-chain MspA protein

While the above experiments demonstrated that single-chain MspA produces functional pores, the purification based on gel extraction is labor intensive and inefficient resulting in low yields of approximately 10 µg per preparation. We estimated that ∼98% of the initial amount of scMspA proteins in the inclusion bodies were lost during this process. One of the main challenges was to separate full-length sc_8_MspA M2 from its many degradation products which are only marginally smaller, i.e. these degradation products may have cleaved linker peptides which appear to be very susceptible to proteolysis (Fig. S6). To address this issue we exploited the fact that both termini of MspA are accessible on the outside of the MspA channel (Fig. 1B). Thus, we added a His_8_-tag at the N-terminus and a Twin-Strep II tag at the C-terminus flanking the *sc*_*8*_*mspA m2* gene (Fig. 3A). We refer to this protein as double-tagged single-chain MspA M2 (sc_8_MspA_dt_ M2). The plasmid pML4170 carrying *sc*_*8*_*mspA*_dt_ *m2* was transformed into *E. coli* BL21(DE3)Omp8 for protein production and purification. Sc_8_MspA_dt_ M2 was purified under denaturing conditions using subsequent Ni(II)- and streptavidin affinity chromatography to isolate full-length protein (Figs. 3B,C, S7). Refolding in the detergent OPOE and a final size exclusion (Fig. S7) yielded 0.2 mg per liter of culture per single preparation of apparently pure protein (Fig. 3 B, C). Importantly, we did not observe contaminating degradation products (Fig. S6) in the refolded protein sample as evident from western blot on Figure 3C. As a quality control we performed denaturing experiments with 80% DMSO. Only octameric MspA M2 dissociated into its monomeric subunits of 20 kDa while sc_8_MspA_dt_ M2 was stable indicating that all eight subunits are covalently linked in the purified refolded protein (Fig. 3D). Overall, the combination of N- and C-terminal affinity tags enabled us to purify full-length scMspA protein with a 20-fold increase in protein yield per single preparation in comparison to the previous gel excision method.

**Fig. 3.**
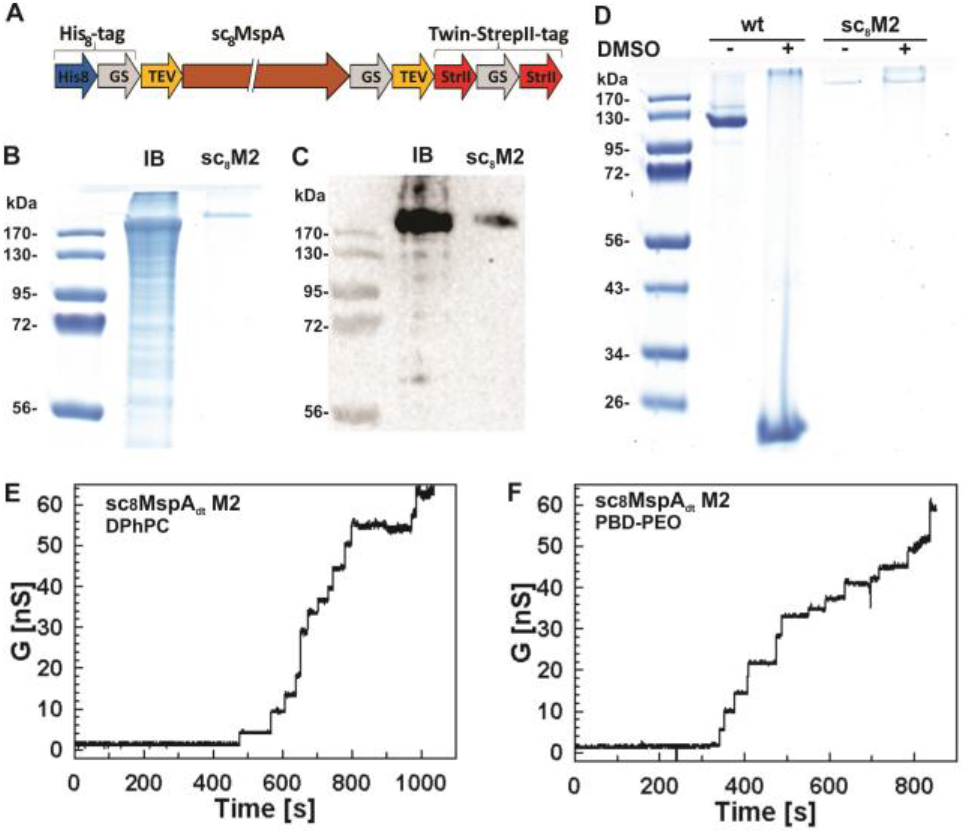
Purification and channel activity of single-chain MspA. **(A)** Scheme of the *scMspA* gene in the expression vector for protein (sc_8_MspA_dt_ M2) purification from E. coli.. Eight *mspA m2* genes (brown arrow) are flanked by two tags: N-terminal His8-tag and C-terminal Twin-StrepII tag. GS – glycine-serin linker; scMspA – single-chain MspA; StrII – StrepII-tag; TEV – TEV protease recognition site. Not to scale. **(B)** Coomassie stained 8% polyacrylamide gel with inclusion bodies (IB) from *E. coli* BL21(DE3) Omp8 and the purified sc_8_MspA_dt_ M2 protein. **(C)** Western blot of the samples shown in B. Proteins were transferred onto PVDF membrane and stained with anti-StrepII-tag HRP-conjugated antibodies. **(D)** Denaturion of MspA M2 and sc_8_MspA_dt_ M2. Coomassie-stained 8% polyacrylamide gel of DMSO treated (+) and untreated (-) samples. Proteins were boiled in 80% (v/v) DMSO to induce denaturing. Note that the 20 kDa band is monomeric subunit of MspA that is visible after DMSO treatment. Lanes: wt – octameric MspA M2; sc_8_M2 – sc_8_MspA_dt_ M2 with eight covalently linked subunits. 3 µg and 1 µg of MspA M2 and sc_8_MspA_dt_ M2, respectively, were loaded. **(E)** Current trace of sc_8_MspA_dt_ M2 in 1 M KCl, 10 mM HEPES, pH 7.4 electrolyte at −10 mV applied potential in diphytanoyl phosphatidylcholine (DPhPC) bilayer with an aperture of approximately 1mm in diameter. The concentration of the protein in the cuvette was 16 ng/ml. **(F)** Current trace of sc_8_MspA_dt_ M2 in 1 M KCl, 10 mM HEPES, pH 7.4 electrolyte at −10 mV applied potential in bi-block polymer (PBD-PEO) bilayer with an aperture of approximately 1 mm in diameter.

### Channel-forming properties of single-chain MspA in large membranes

To examine whether sc_8_MspA_dt_ M2 forms functional channels we performed lipid bilayer experiments in a Montal-Mueller setup using cuvettes with an aperture diameter of 1 mm. As expected, refolded sc_8_MspA_dt_ M2 had channel-forming activity in DPhPC membranes as shown by the stepwise current increase after addition of the protein (Fig. 3E). Analysis of 432 insertions from 8 different membranes showed predominant conductance peaks of 1.3 nS, 1.9 ns, and 3.0 nS (Fig. S4E). Notably, sc_8_MspA_dt_ M2 had a broader conductance distribution than sc_8_MspA PN1 (Fig. S5F). These differences may be the result of different purification methods, the absence of the P97F mutation near the constriction zone of sc_8_MspA_dt_ M2 compared to sc_8_MspA PN1, and/or the presence of purification tags on the termini of sc_8_MspA_dt_ M2.

We recently demonstrated that octameric MspA M2 forms channels in bilayers composed of poly(1,2-butadiene)-b-poly(ethylene oxide) (PBD-PEO) polymer (10). PBD-PEO bilayers have a two-to three-fold increased lifetime and are more robust towards chemicals and high voltages than membranes made from biological lipids, and can be used in nanopore sequencing experiments (10). Thus, we also examined the channel properties of scMspA insertions in PBD-PEO bilayers. Similarly to DPhPC bilayers, we observed insertions of sc_8_MspA_dt_ M2 into PBD-PEO bilayers (Fig. 3F). Analysis of 207 insertions from 28 membranes showed broad distribution of single-channel conductances (Fig. S4F) in PBD-PEO bilayers similar to those obtained for sc_8_MspA_dt_ M2 in DPhPC bilayers. It should be noted that insertion frequency of sc_8_MspA_dt_ M2 was reduced in PBD-PEO bilayers in comparison to DPhPC membranes. Taken together, these experiments showed that sc_8_MspA_dt_ M2 forms functional channels in both DPhPC and PBD-PEO bilayers.

### Hairpin translocation through the single-chain MspA pore

To examine whether the nucleotide recognition capability of MspA is preserved in scMspA, we used sc_8_MspA_dt_ M2 in DNA hairpins experiments as shown previously (12). This assay is based on distinct residual currents when the single-stranded homopolymer tail is located inside the constriction zone (Table S4), while the double-stranded region of the DNA hairpins is stalled in the lumen of the MspA pore and temporarily prevents translocation (Fig. 4A). Single sc_8_MspA_dt_ M2 pores were inserted into poly(1,2-butadiene)-b-poly(ethylene oxide) (PBD_11_–PEO_8_) block-copolymer bilayer membranes with a diameter of 100 µm and bathed in a buffer containing 10 mM Tris (pH 7.5), 1 M KCl and 2 M guanidinium chloride (GdmCl). Based on previous data (10) we used GdmCl in the electrolyte solution because it induces a semi-melting state of DNA hairpins enabling smooth DNA passage through the nanopore at low applied voltage resulting in increased DNA capture rate and decreased trapping time.

**Fig. 4.**
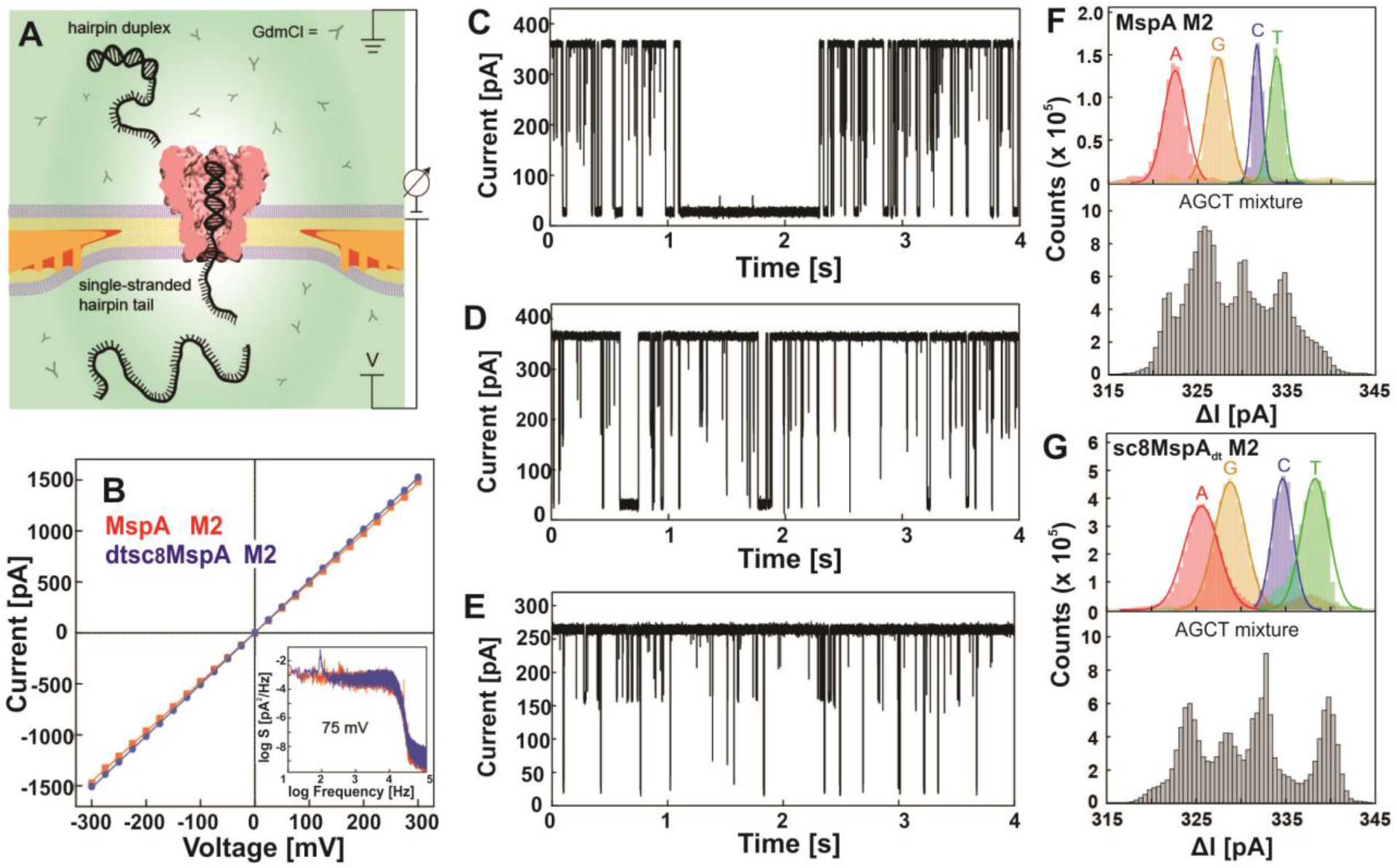
DNA recognition by single-chain MspA. DNA hairpin translocation experiments with MspA M2 and sc_8_MspA_dt_ M2 were performed in free-standing PBD-PEO polymer bilayers supported by wedge-on-pilar 100 µm aperture at 75 mV applied voltage in 1M KCl, 2 M CgmCl, 10 mM Tris, pH 7.5 by patch-clamp amplifier. **(A)** A free-standing PBD-PEO polymer bilayer membrane is supported by wedge-on-pillar aperture. A single mutant MspA nanopore inserted is shown in the buffer with 2 M GdmCl. A patch-clamp amplifier is connected by Ag/AgCl to measure the ion current through the *cis* (top) and *trans* (bottom) chambers. **(B)** I-V curves of sc_8_MspA_dt_ M2 and MspA M2 from − 300 to 300 mV with the power spectral density plots at 75 mV shown, in 1 M KCl, 2 M GdmCl, 10 mM Tris, pH 7.5 electrolyte. **(C, D)** Continuous current vs. time traces for MspA M2 (C) and sc_8_MspA_dt_ M2 (D) in 1M KCl, 2 M GdmCl, 10 mM Tris, pH 7.5 buffer with 75 mV voltage applied, showing the translocation events caused by DNA hairpin poly-dT going through the nanopore. **(E)** Continuous current vs. time traces for sc_8_MspA_dt_ M2 in 1M KCl, 10 mM Tris, pH 7.5 buffer with 140 mV voltage applied. **(F, G)** Histograms of current change (ΔI) caused by the blockade of individual DNA hairpins(top panel) and DNA hairpin mixture (bottom panel) for MspA M2 (F) and sc_8_MspA_dt_ M2 (G) in 1 M KCl, 2 M GdmCl, 10 mM Tris, pH 7.5 electrolyte and 75 mV voltage. 0.3 µM of each DNA hairpin (poly-dA, poly-dC, poly-dG and poly-dT) were added.

The current-voltage curves and power spectral density plots reveal that the conductance and trace noise are almost identical for sc_8_MspA_dt_ M2 and octameric MspA M2 (Fig. 4B). Both proteins exhibited gating when negative voltage was applied (Fig. S8A). Since the gating conductance varies for the same MspA pore even at the same voltage, we used the ungated current values for the IV curves resulting in identical symmetric IV curves for both sc_8_MspA_dt_ M2 M2 and MspA M2 (Fig. 4B). Uncorrected IV curves for the same raw data are shown in Figure S8B. Interestingly, the protein concentration required to observe channel insertions was approximately 60-fold higher for sc_8_MspA_dt_ M2 (4.2 nM) than for octameric MspA M2 (0.063 nM). Consistent with previous results in large aperture bilayers (Fig. S4F), a wide conductance distribution was observed for individual sc_8_MspA_dt_ M2 channel ranging from 1 nS to 5.5 nS. However, only the pore with conductance of around 4.9 nS resulted in clean and deep DNA translocation signals. This appears to be a general property of the MspA pore as populations of pores with different conductance levels were previously observed in hairpin DNA experiments with octameric MspA isolated from *M. smegmatis* (25).

The current traces after addition of the poly-dT hairpin to sc_8_MspA_dt_ M2 and MspA M2 show current blockades resulting from translocation of the DNA hairpin through the pore in GdmCl-containing buffer (Fig. 4C, D). Similar current blockades were observed with the poly-dT hairpin and sc_8_MspA_dt_ M2 when no GdmCl was present in the electrolyte buffer (Fig. 4E). However, a higher voltage of 140 mV instead of 75 mV was necessary to overcome the energy barrier for DNA translocation in accordance with previous hairpin experiments with octameric MspA where 140 to 180 mV was used for hairpin DNA translocation (12). Our data further indicate that GdmCl improves both capture and translocation of hairpin DNA through the pore.

### Nucleotide recognition by single-chain MspA

To examine nucleotide recognition by sc_8_MspA_dt_ M2 we used DNA hairpins with a duplex region of 14 nucleotides and a homopolymer tail of 50 nt (hp14-dT50 (poly-dT), hp14-dA50 (poly-dA), hp14-dC27 (poly-dC), and hp14-dG3dA47 (poly-dG) (Table S3, Fig. S9). It should be noted that we used only three dG nucleotides in the “poly”dG hairpin tail in otherwise dA background to avoid G-tetrad formation as in our previous experiments (12). Histograms of the current change ΔI (current I in the presence of a hairpin minus the baseline current level I_0_) are shown for the four individual DNA hairpins and mixture of four hairpins with octameric MspA M2 (Fig. 4F) and sc_8_MspA_dt_ M2 (Fig. 4G), respectively. Fitted Gaussians for each individual hairpin with a homopolymer tail were well-resolved and separated, and looked almost identical for both MspA M2 and sc_8_MspA_dt_ M2 (Fig. 4F, 4G, top panel). The partial overlap between purines was more pronounced for sc_8_MspA_dt_ M2, nonetheless, individual peaks for poly-dA and poly-dG were still distinguished (Fig. 4G, top panels). Remarkably, the histograms of DNA hairpin mixture clearly show four separate peaks representing the four different nucleotides, which match the current blockades for individual hairpin peaks (Fig. 4F, 4G, bottom panels). In this experiment sc_8_MspA_dt_ M2 had better resolving properties than octameric MspA M2. In conclusion, our DNA hairpin experiments showed that sc_8_MspA_dt_ M2 and octameric MspA M2 have very similar capabilities in distinguishing all four DNA nucleotides. These results indicate that covalently linking of all eight MspA subunits preserves the nucleotide recognition properties of the MspA pore and open many avenues to tailor the channel properties of the MspA pore for specific sensing applications.

## DISCUSSION

### Properties and potential applications of single-chain MspA nanopores

In this study we showed that it is feasible to convert multimeric MspA into a functional monomeric nanopore with identical electrophysiological properties. This achievement is a major improvement over previous approaches based on self-assembly of wt and mutated subunits, which required the physical separation of pores with different subunit combinations lowering the yield dramatically, and suffered from different permutations with identical physical properties (3). By contrast, single-chain MspA can be produced as a full-length protein with many asymmetric mutations opening numerous avenues to improve the performance of MspA in DNA sequencing. For example: (i) Distinct amino acids in the MspA constriction zone will enable different chemical interactions with the DNA nucleobases. Previously, mutations that enable DNA translocation were present in all eight MspA monomers (11), making asymmetric interactions with DNA impossible. Asymmetric mutations open numerous avenues to increase the specificity of the current blockade for each nucleotide and to increase the dwell time in the construction zone. Both of these effects are likely to reduce the contribution of neighboring nucleotides to the current blockade, which is currently 4–5 nucleotides (30), and concomitantly reduce the high raw data error rates in nanopore sequencing (31). (ii) Single-chain MspA can be used to slightly alter the diameter of the pore to modulate the interactions between amino acids in the constriction zone and nucleotides. This can be achieved changing the subunit stoichiometry as shown in this study (Fig. 2). In a simple geometrical model adding or removing a subunit alters the diameter of the MspA pore by 1/8^th^, which is equivalent to approximately 1.5 Å, a reasonable range for significantly changing chemical interactions. Both the hexameric and heptameric pores are functional demonstrating that this approach is feasible. (iii) Motor proteins such as DNA polymerases employed to control the translocation rate of DNA through the pore have been instrumental for sequencing (13) and have become standard in nanopore sequencing devices (32). However, the long distance between the motor proteins and the MspA constriction zone descreases the positional precision and increases error due to thermal motions of the flexible DNA strand (33) and the motor protein. Random positioning of the motor protein on top of the MspA pore might enhance this error. A single cysteine in one of the surface loops of single-chain MspA would enable the covalent attachment of the motor protein, eliminating the random positioning of motor and assembly with the MspA pore, and thereby reducing the system variability. (iv) It might be possible to create a specific path for single-stranded DNA inside of the MspA pore, thereby decreasing the translocation velocity and eliminating the need for a motor protein althogether. While the possibilities to tailor the single-chain MspA pore for DNA sequencing by combining asymmetric mutations are almost endless, it would require an efficient assay to test these combiniations. However, the currently used lipid bilayer systems are not high-throughput. By contrast, the single molecule measurements are tedious and usually require the analysis of several pores before a pore suitable for sequencing is identified. Hence, the recent developments of computer modelling for structure predictions (34,35) and virtual drug screening (36) could provide a valuable alternative to test and identify beneficial combinations of mutations of the single-chain MspA pore *in silico*.

### Heterogeneous channel conductances of MspA pores

Our study showed a wide variety of channel conductances for our single-chain MspA constructs ranging from 0.5 nS to 4.5 nS for sc_8_MspA M2 without tags (Fig. S4D). This heterogeneity makes identification of single-chain MspA pores suitable for sequencing more difficult. However, this is not a consequence of the covalent linkers between the subunits of single-chain MspA as the channel conductances of the unmodified octameric MspA pore isolated from *M. smegmatis* also vary between 2 and 5 nS (Fig. S4A). These measurements are consistent with early measurements (6,37) indicating that the conductance variability is an intrinsic property of the MspA pore. The mutations introduced in MspA M1 and MspA M2 to enable DNA translocation (11) appear to increase the intrinsic conductance variability but shift the major conductance peak from 4.5 - 5 nS to app. 1.5 nS (Figs. S4B, C), indicating a significant contribution of the constriction zone to the overall channel conductance of MspA. It is plausible that neutralizing the constriction zone by mutating the negatively charged aspartates D90 and D91 to asparagines leads to a smaller and more flexible constriction zone due to the lack of electrostatic repulsion explaining both the shift to smaller channel conductances and their wider range. While we cannot exclude that the harsh procedures used to extract MspA from *M. smegmatis* such as organic solvents (6) or boiling in detergents (27) contribute to pore heterogeneity, it is striking that the recombinant single-chain MspA M2 pores, which were isolated from *E. coli*, showed a similar conductance profile as octameric MspA M2 isolated from *M. smegmatis*. Thus, it appears that pore conductance heterogeneity is largely an intrinsic property of MspA, driven mainly by the properties of the constriction zone. The hypothesis that the flexible constriction zone of the MspA variants M1 and M2 used in DNA sequencing experiments and in the corresponding single-chain variant sc_8_MspA M2 is major driver of the conductance heterogeneity of these pores is supported by an additional experimental observation. The MspA PN1 variant with a phenylalanine 97 at the tip of the loop 6, which connects the constriction zone with the subsequent β-strand (38,39), has a much smaller conductance distribution (Fig. S5A), indicating that putative hydrophobic interactions of neighboring phenylalanines stabilize the constriction zone. Conductance heterogeneity is common in general diffusion pores from other bacteria. For example, OmpF, the main porin of *E. coli*, and porins of *Salmonella* and *Pseudomonas aeruginosa* show a broad range of channel conductances (40-43). By contrast, the channel conductances are much better defined for specific porins, which have specific substrate binding sites, and the much smaller ion channels, which are located in the cytoplasmic membrane. For example, LamB, a maltodextrine-specific porin of *E. coli* (44-46), OprO, a phosphate-specific porin of *P*.*aeruginosa* (47), and the bacterial amyloid secretion channel CsgG (48) have narrow conductance distributions.

### Single-chain MspA pores with different subunits stoichiometries

In this study we constructed scMspA pores with different subunit stoichiometries to demonstrate the feasibility of changing the channel diameter, an important feat for nanopore sensing applications. All four single-chain MspA constructs with less than eight subunits formed functional pore proteins despite the likely limited tolerance of the MspA structure, in particular of the β-barrel, for expansions or reductions of the number of subunits. This is probably due to the self-assembly of the scMspA constructs with three and five subunits considering that the effiency of the assembly of covalently linked subunits to a functional pore probably decreases with larger deviations from the octameric pore, which was found previously to be the most dominant form in a self-assembly process with the purified MspA monomer (38). Interestingly, the sc_8_MspA PN1 pore had a much broader channel distribution than octameric MspA PN1 indicating that the purification and refolding procedure of recombinant scMspA introduced channel heterogeneity, which is not observed for octameric MspA PN1 purified from *M. smegmatis*. Thus, we expected that sc_3_MspA PN1 assembles mostly to a hexameric structure. This is consistent with the conductance profile obtained for sc_3_MspA PN1, which resembles that of sc_6_MspA PN1 (Fig. S5B, D). The few additional larger channels for sc_3_MspA PN1 might have been from a nonameric MspA pore. The conductance distribution of sc_5_MspA PN1 is shifted to larger values compared to sc_8_MspA PN1 indicating the presence of mainly decameric MspA pores. The conductance profile of sc_6_MspA PN1 is similar to that of sc_8_MspA PN1, while the conductances of sc_7_MspA are shifted to larger values indicating assembly to larger pores (Fig. S5). Structural analysis of the single-chain variants by electron microscopy will be necessary to obtain quantitative results about different scMspA assembly forms for each subunit composition and exact constriction zone diameters of the scMspA pores.

## Conclusions

This study demonstrated the feasibility of producing single-chain MspA pores and their use in DNA sequencing. The unique adaptability of the single-chain MspA pore as a biosensor by making asymmetric mutations has enormous potential to improve the DNA sequencing capability of MspA and for many other applications such as detection of RNA (49), proteins (50) and small molecules (51). The single-chain concept might be applicable to other oligomeric protein pores such as CsgG(52), ClyA (53), FraC(54) and α-HL(55), depending on the proximity of the C-terminus of one monomer to the N-terminus of the next monomer in the pore structure.

## ACKNOWLEDGEMENTS

This work was supported by the National Institutes of Health grant R21 HG010543 to M.N. The funding source had no involvement in preparation of the article, study design, in the collection, analysis and interpretation of data; in the writing of the report; and in the decision to submit the article for publication.

